# Novel immune cell subtypes linked to survival among African American women with triple-negative breast cancer

**DOI:** 10.1101/325951

**Authors:** Kristen S. Purrington, Andreana N. Holowatyj, Michele L. Cote, Ann G. Schwartz, Rahman Chaudhry, Rouba Ali-Fehmi, Gregory Dyson, Justin Colacino, Julie Boerner, Sudeshna Bandyopadhyay

**Affiliations:** Department of Oncology, Wayne State University School of Medicine, Detroit, Michigan; Department of Pathology, Wayne State University School of Medicine, Detroit, Michigan; Department of Environmental Health Sciences, School of Public Health, University of Michigan, Ann Arbor, Michigan; Population Studies and Disparities Research Program, Barbara Ann Karmanos Cancer Institute, Detroit, Michigan; Molecular Therapeutics Program, Barbara Ann Karmanos Cancer Institute, Detroit, Michigan; Tumor Biology and Microenvironment Program, Barbara Ann Karmanos Cancer Institute, Detroit, Michigan

**Keywords:** tumor-infiltrating lymphocytes, gene expression, pathology, survival, immune, inflammation

## Abstract

Triple negative breast cancer (TNBC) is an aggressive disease that is twice as likely to be diagnosed in African American (AA) women compared to white women, with poor clinical outcomes. Tumor infiltrating lymphocytes (TILs) are associated with improved survival for TNBC, but the relevance of TILs and immune cell subtypes to survival in AA women with TNBC is unknown. We evaluated histopathologic TIL counts and molecular characteristics among 60 AA women diagnosed with TNBC with linkage to clinical outcomes using data from the Metropolitan Detroit Cancer Surveillance System. We utilized whole genome expression profiling of TN tumors and cell type deconvolution analysis to evaluate the underlying mechanisms and immune cell subtypes associated with survival patterns in the context of TILs. TILs were significantly associated with improved survival [1-10% Hazard Ratio (HR)=0.32, 95% Confidence Interval (CI) 0.12-0.90, p=0.031; >10% HR=0.18, 95% CI 0.05-0.67, 9.9×10^−3^]. 524 transcripts (326 coding, 198 non-coding) were associated with TIL levels, 34 of which were associated with both TILs and survival (p<0.05). While only naïve B cells were associated with survival when considering individual cell types [Median HR=2.43, 95% CI 1.07-5.55, p=0.035], increased naïve B cells, plasma cells, and activated NK cells, and decreased resting mast cells, M1 macrophages, and monocytes were associated with transcripts that predicted worse survival. These data provide evidence for novel roles for these immune cells types in TNBC, and further studies are needed to validate these findings and identify determinants of patterns of immune response in TNBC relevant to the AA population.

**Summary:** We found that increased naïve B cells, plasma cells, and activated natural killer cells, and decreased resting mast cells, M1 macrophages, and monocytes were associated with expression biomarkers of worse survival among African American women with triple negative breast cancer.

## INTRODUCTION

Triple negative breast cancer (TNBC), defined by low or no expression of estrogen receptor (ER), progesterone receptor (PR), and human epidermal growth factor receptor-2 (HER2), is an aggressive disease that is twice as likely to be diagnosed in Black/African American women compared to white/European American women (1). Overall clinical outcomes for TNBC are poor yet heterogeneous: peak death and recurrence rates (~30%) occur within the first three years following treatment, but survival rates for the women who live beyond this timeframe are similar to those of hormone receptor positive breast cancers (2). Poor TNBC survival is a large contributor to the racial disparity in overall breast cancer outcomes since African American women are at higher risk for this subtype, where African American women are 1.4-fold more likely to die from overall breast cancer than European American women (3). Several studies further suggest that African American women with TNBC specifically experience poorer clinical outcomes compared to European American women (4–9), although this relationship is less clear.

Tumor infiltrating lymphocytes (TILs) are a well-established, favorable prognostic factor for overall breast cancer, including TNBC, that may in part account for the observed heterogeneity in prognosis among women with TNBC (10–12). These mononuclear immune cells, representing a heterogeneous population of T cells, B cells, natural killer (NK) cells, and others, infiltrate tumor tissue into either the desmoplastic stroma (stromal TILs, sTILs) and/or into tumor cell clusters (intratumoral TILs, iTILs), and are representative of a patient’s immunological response to the tumor (13). TILs have recently been described as most prevalent in TNBC compared to other breast cancer subtypes, where ~20% of tumors exhibit high levels of TILs (>50% lymphocytic infiltrate), which seems to synergize with chemotherapeutic agents to provide a 30% reduction in risk of death (10–12,14–16). However, 10-20% of TN tumors are estimated to have no immune infiltrate, representing a subpopulation of TNBC patients that may benefit from strategies to enhance immunity (10). This would be particularly impactful for African American women with TNBC, for whom there is evidence that TNBC clinical outcomes are worse than European American women (7–9). Indeed, there is evidence that substantial racial differences in tumor immunobiology and microenvironment exist and may contribute to racial differences in survival, although this has not been evaluated in TNBC to our knowledge (17–20).

Although histopathologic TIL counts provide strong prognostic information, the relative contributions of the underlying cell types represented by this summary measure to survival remain unknown. To date, studies of TILs in TNBC have overwhelmingly focused on the contribution of CD8+ T cells to this response, including CD4+ T helper 1 (T_H_1) cells that facilitate direct lysing of tumor cells via cytokines and FOXP3+ T regulatory (T_REG_) cells that dampen CD8+ immune response (10,21–23). However, these approaches capture an incomplete picture of the complex, multifactorial immune response. Two recent studies utilizing publically available expression microarray data examined the association between 22 immune cell subtypes and clinical outcomes in breast cancers using bioinformatic cell type deconvolution (24,25). The one study able to stratify by ER, PR, and HER2 status found that higher fractions of neutrophils, resting mast cells, M2 macrophages, resting NK cells, and resting CD4+ memory T cells were associated with worse overall survival, and better overall survival was associated with higher fractions of plasma and naïve B cells among TNBC (25). However, no racial information was utilized in this analysis, so the relevance of these immune subtypes for African American women is unclear. Here, we evaluate histopathologic TIL counts and molecular characteristics of TNBC lesions among a cohort of African American women. Using gene expression profiles and cell type deconvolution, we describe potential mechanisms and immune cell populations that mediate the association between TILs and survival.

## METHODS

### Study population and follow-up

Tumor block identification, retrieval, and linkage with clinical information were performed by the Karmanos Cancer Institute Biobanking and Correlative Sciences Core and the Epidemiology Research Core. African American women who had surgery for clinically confirmed primary invasive ER/PR/HER2-negative breast cancer from 2004 through 2015 at the Karmanos Cancer Center in Detroit, Michigan were identified through the Metropolitan Detroit Cancer Surveillance System (MDCSS), a founding member of the National Cancer Institute’s Surveillance, Epidemiology, and End Results (SEER) Program. ER- and PR-negative tumors were identified by SEER records, and HER2-negative status was determined through a combination of SEER (diagnosed 2010 or later) and pathology records (diagnosed prior to 2010). A total of 239 eligible women were identified; tumor samples were obtained for sixty of these patients for expression profiling. Clinical, treatment, and outcomes data were obtained via linkage with the SEER registry, including stage, grade, laterality, age at diagnosis, ER/PR/HER2 status, surgery type, systemic therapy type, radiation, sequence of surgery and systemic therapy (among women who had chemotherapy), vital status at last contact, cause of death, and date of last contact. Menopausal status was obtained from medical record review. This study was approved for exemption by Wayne State University Institutional Review Board.

### Histopathology and gene expression profiling

Slides were cut from formalin-fixed paraffin-embedded (FFPE) blocks and stained with hematoxylin and eosin (H&E) in the Biobanking and Correlative Sciences Core at the Karmanos Cancer Institute. Tissue curls were generated from four 10mm unstained slides corresponding to the tumor area and collected in DNAse/RNAse free microcentrifuge tubes. Total RNA was extracted using the QIASymphony Automated system (Qiagen, Germany) according to the manufacturer protocol. Gene expression profiling was performed using Affymetrix Human Gene ST 2.0 arrays after amplification of RNA using the Affymetrix WT Pico Kit (Santa Clara, CA). Raw probe intensity data was exported for statistical analysis.

### TIL scoring

H&E slides were reviewed by a single breast cancer pathologist (S.B.). TILs were scored using methods outlined by the International TILs Working Groups (26) for both stromal (sTIL) and intratumoral (iTIL) tumor-infiltrating lymphocytes separately. All mononuclear cells, including lymphocytes and plasma cells, and excluding polymorphonuclear leukocytes and granulocytes, inside the boundaries of the invasive tumor were included. sTILs were defined as those within the stroma but not in contact with tumor cells, and were scored as present versus absent. iTILs were scored using the following categories: 0%, 1-10%, 11-50%, >50%.

### Bioinformatics

Pathway analyses were conducted using Ingenuity^®^ Pathway Analysis (IPA) software (Qiagen, Hilden, Germany). Enrichment for canonical pathways was evaluated, where we considered pathways with p-values <0.001 and at least five molecules from our gene list were present. Cell type deconvolution was performed as implemented in the CIBERSORT program (27). CIBERSORT is a method for estimating the cellular composition of complex tissues from gene expression profiles, including solid tumors. We utilized the “LM22” signature gene file as the input reference gene signature, representing the validated 547-gene expression profile of 22 mature human hematopoietic populations and activation states including the following: naïve B cells, memory B cells, plasma cells, CD8+ T cells, naïve CD4+ T cells, CD4+ memory T cells, activated CD4+ memory T cells, follicular helper T cells, regulatory T cells, γδ T cells, resting NK cells, activated NK cells, monocytes, M0 macrophages, M1 macrophages, M2 macrophages, resting dendritic cells, activated dendritic cells, resting mast cells, activated mast cells, eosinophils, and neutrophils. CIBERSORT was run using default options and 100 permutations.

### Statistical analysis

Due to the small number of tumors containing >50% iTILs (n=9),we collapsed the variable to categories of 0%, 1-10%, and >10% for all analyses. Chi-squared tests, Fisher’s exact tests, and t-tests were used to evaluate associations between iTILs and clinical/pathologic characteristics. Associations between both iTILs (coded 0, 1, 2 corresponding to 0%, 1-10%, >10%) and sTILs (present vs. absent) considered as categorical variables and overall survival were evaluated in Cox proportional hazards (CoxPH) models as implemented in the “survival” package in R (https://cran.r-project.org/), adjusting for stage and age at diagnosis. Subset analyses among women with known chemotherapy sequence (adjuvant vs. neoadjuvant) were performed to evaluate the effect of including treatment type and chemotherapy sequence as covariates in survival models. Raw probe intensity data were normalized as implemented by the “rma” function in R to perform background subtraction, quantile normalization, and summarization of probe sets using median-polish, resulting in 44,629 transcript probes for analysis (27,016 coding, 17,613 non-coding). Given that sTILs were present in more than 85% of tumors and the previously reported associations between iTILs and survival (28–32), we focused all survival and genetic analyses on iTILs. Given that categorical iTIL associations with survival were consistent with a log-linear relationship between iTIL category and survival, subsequent analyses were conducted using iTILs as an ordinal variable. We evaluated associations between iTILs and each of the 44,629 transcripts using linear regression. Transcripts were selected for subsequent analyses using a liberal significance threshold of p<0.01 (n=524), and pathway analysis was utilized to demonstrate that these associations represented significant enrichment of biologically relevant immune pathways. These transcripts were evaluated as predictors of survival in CoxPH models adjusting for age, stage, and iTIL level. We then formally evaluated these transcripts as causal mediators of the association between iTILs and survival using the “mediation” package in R, which provided effect estimates and p-values for the direct effect of iTILs on survival, the mediation effect of the transcript, and the proportion of the iTIL effect that is mediated by the transcript. This package allowed for parametric modelling, such that we individually fit 1) gene expression predicted by iTIL level via linear regression and 2) the survival model with iTILs, transcript probe, and age and stage at diagnosis as predictors using the “survreg” function. Given that transcripts were evaluated on the basis of a priori evidence of association with iTILs, mediation was considered statistically significant if the mediation and direct effect p-values for the transcript probes were both <0.05. Cell type proportions from CIBERSORT analysis were evaluated dichotomized by their median in CoxPH models adjusting for iTILs, stage, and age at diagnosis for associations with survival. Cell type proportions were also evaluated for correlations with the expression of 34 transcripts of interest in linear regression models.

## RESULTS

### TNBC cohort

Sixty African American women from metropolitan Detroit with primary invasive triplenegative breast cancer for whom tumor blocks were available comprise our study cohort. A total of 75% of cases were pathologically detected to have intratumoral tumor-infiltrating lymphocytes (iTILs) present and more than 85% had stromal TILs present (Table 1, **Figure S1**). Mean age at diagnosis was significantly associated with iTILs, where women with any iTILs present were diagnosed with TNBC nearly 10 years younger than women with no iTILs (p=6.7×10^−3^). Similarly, menopausal status was significantly associated with iTIL levels, where nearly half of women with any iTILs were pre-menopausal compared to less than 10% of women with no iTILs (p=0.038). Among women in our cohort, there were no significant differences in stage at diagnosis and laterality by level of iTILs detected in the tumors (Table 1). Among those with known type of surgical therapy, 58.6% of cases underwent surgical mastectomy for primary TNBC treatment. The majority of patients received chemotherapy treatment (>90%), while fewer patients underwent radiation therapy for their breast cancer (>70%). Although chemotherapy data from SEER are incomplete, sub-set analyses among those with known chemotherapy type allowed us to evaluate potential differences in associations for those with neoadjuvant versus adjuvant chemotherapy. There was no association between therapy type or sequence of systemic therapy and surgery and iTILs.

**Table 1.** Clinical, pathological, and demographic characteristics of African American women TNBC by iTIL category.

| Intratumoral Tumor-Infiltrating Lymphocytes (iTILs) |  |  |  |  |  |  |  |
| --- | --- | --- | --- | --- | --- | --- | --- |
| Characteristic | 0% |  | 1-10% |  | >10% |  | *p-value* ** |
|  | *N* | % | *N* | % | *N* | % |  |
|  | 15 |  | 29 |  | 16 |  |  |
| **Age at Diagnosis** |  |  |  |  |  |  | 0.11 |
| ≤55 years | 6 | 40.0% | 21 | 72.4% | 10 | 62.5% |  |
| 56+ years | 9 | 60.0% | 8 | 27.6% | 6 | 37.5% |  |
| Mean (std) | 58.2 | (12.3) | 48.3 | (9.5) | 48.6 | (15.3) | 6.7x10<sup>-3</sup> |
| **Menopausal status** |  |  |  |  |  |  | 0.038 |
| Pre-menopausal | * | <10% | 11 | 39.3% | 8 | 50.0% |  |
| Post-menopausal | * | >90% | 17 | 60.7% | 8 | 50.0% |  |
| **Stromal TiLs (sTiL)** |  |  |  |  |  |  | 0.010 |
| None | 5 | 33.3% | 1 | 3.6% | 1 | 6.3% |  |
| Yes | 10 | 66.7% | 27 | 96.4% | 15 | 93.7% |  |
| **Laterality** |  |  |  |  |  |  | 0.98 |
| Right | 9 | 60.0% | 17 | 58.6% | 9 | 56.3% |  |
| Left | 6 | 40.0% | 12 | 41.4% | 7 | 43.8% |  |
| **Stage** |  |  |  |  |  |  | 0.28 |
| Localized | 10 | 66.7% | 14 | 48.3% | 7 | 43.8% |  |
| Regional/Distant | 5 | 33.3% | 15 | 51.7% | 9 | 56.3% |  |
| **Surgical Therapy**<sup>†</sup> |  |  |  |  |  |  | 0.76 |
| Breast-conserving surgery | 5 | 33.3% | 12 | 41.4% | 7 | 43.8% |  |
| Mastectomy | 10 | 66.7% | 15 | 51.7% | 9 | 56.3% |  |
| **Chemotherapy**<sup>‡</sup> |  |  |  |  |  |  | 0.72 |
| No | * | < 15% | * | < 10% | * | < 7% |  |
| Yes | * | ≥ 85% | * | ≥ 90% | * | ≥ 93% |  |
| **Chemotherapy sequence**<sup>‡</sup> |  |  |  |  |  |  | 0.99 |
| Neoadjuvant chemotherapy | * | <45% | 9 | 47.4% | 7 | 53.8% |  |
| Adjuvant chemotherapy | * | >55% | 10 | 52.6% | 6 | 46.2% |  |
| **Radiation Therapy** |  |  |  |  |  |  | 0.47 |
| No | * | < 30% | 6 | 20.7% | 6 | 37.5% |  |
| Yes | * | ≥ 70% | 23 | 79.3% | 10 | 62.5% |  |
| **Status** |  |  |  |  |  |  | 0.062 |
| Alive | 5 | 33.3% | 17 | 58.6% | * | ≥ 75% |  |
| Dead | 10 | 66.7% | 12 | 41.4% | * | < 25% |  |

\* Values were censored due to small sample size.
\*\*p-value excludes unknown values.
† Two patients had unknown surgical therapy type.
‡ Among patients with chemotherapy data

### TILs & TNBC survival

A total of 26 (43.3%) women died over a median follow-up time of 62 months [Interquartile range (IQR): 21.8-85.5 months] for the total cohort, of which 23 were deaths due to breast cancer (Figure 1a). The median survival time for those who died was 20 months [IQR: 15.0-30.8 months], where 76.6% of women died within 3 years of their diagnosis. Consistent with published findings (28,29,32–34), the presence of iTILs was significantly associated with survival, where each increase in iTIL level was associated with a 60% reduction in the hazard of death adjusting for age and stage at diagnosis [ordinal Hazard Ratio (HR)=0.41, 95% Confidence Interval (CI) 0.21-0.82, p=0.014; 1-10% Hazard Ratio (HR)=0.32, 95% Confidence Interval (CI) 0.12-0.090, p=0.031; >10% HR=0.18, 95% CI 0.05-0.67, 9.9×10^−3^] (Table 2, Figure 1b). Given that these associations were consistent with a log-linear association between iTIL category and survival [HR=0.41, 95% CI 0.21-0.81, p=0.010], subsequent analyses were conducted using iTILs as an ordinal variable. Stromal TILs were not significantly associated with survival adjusting for stage and age at diagnosis [HR=1.22, 95% CI 0.29-5.19, p=0.078], partially accounted for by the fact that sTILs were present in the vast majority of tumors (>85%). Results were nearly identical for breast cancer specific survival given that nearly 90% of deaths were due to breast cancer [1-10% iTILs HR=0.27, 95% Confidence Interval (CI) 0.09-0.78, p=0.016; >10% iTILs HR=0.13, 95% CI 0.03-0.53, p=4.4×10^−3^] (**Table S1**, **Figure S2**). Although systemic therapy and surgery sequence were not associated with TILs, we also evaluated the impact of neoadjuvant chemotherapy (NAC) compared to adjuvant chemotherapy on the association between TILs and survival among women with known chemotherapy type. In a multivariable model, NAC was associated with increased risk of death; however, timing of chemotherapy did not impact the magnitude of association between iTILs and overall or breast cancer specific survival (Table 2, **Table S1**), regardless of the inclusion of surgery type or radiation treatment. We thus utilized Model 1, with iTILs, age at diagnosis, and stage as predictors, for all subsequent survival analyses.

**Figure 1.**
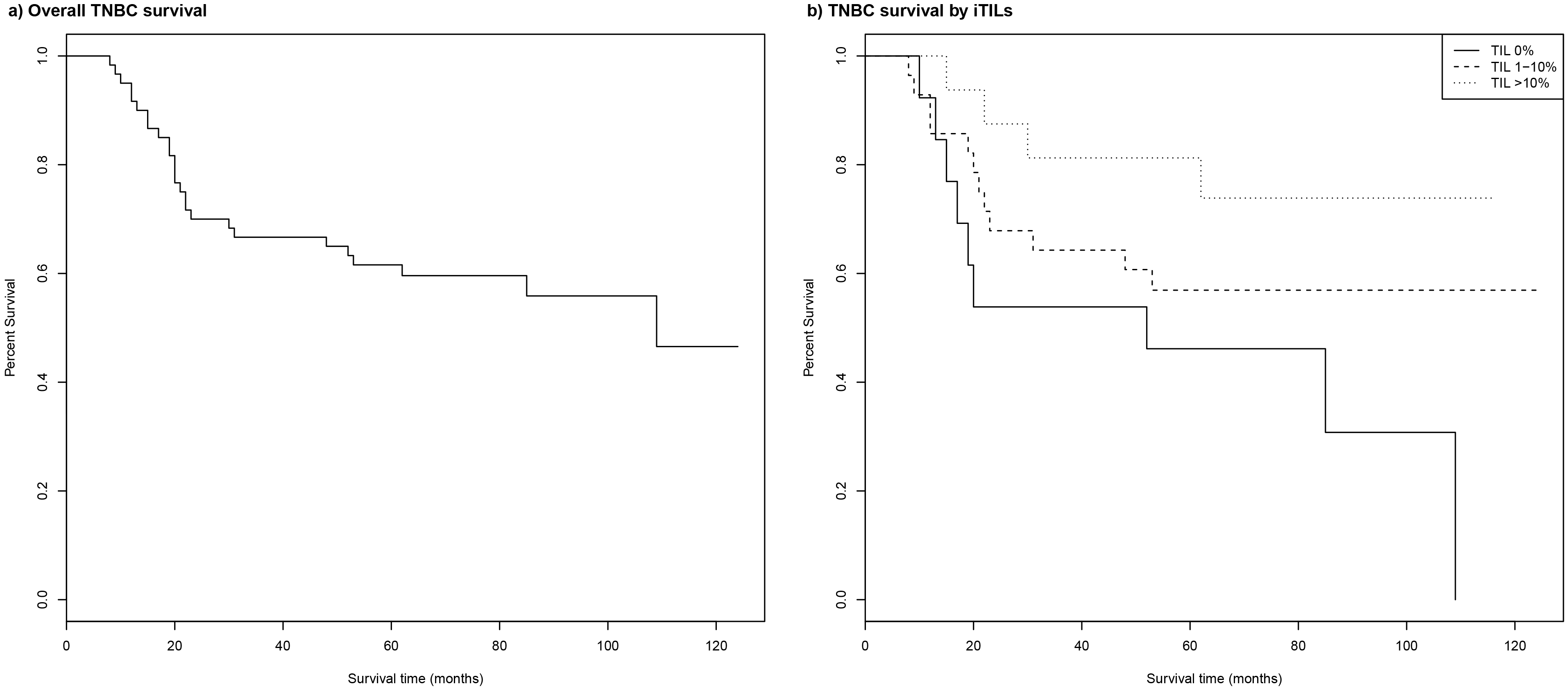
Overall survival among 60 African American women diagnosed with triple-negative breast cancer (TNBC) Kaplan-Meier plots for (a) overall survival and (b) overall survival by TIL level are shown.(a) Solid line represents Kaplan-Meier curve from the total cohort. Dotted lines represent 95% confidence limits. (b) Solid lines represent Kaplan-Meier curves by TIL level: 0%=solid, 1-10%=dashed, >10%=dotted.

**Table 2.** Multivariable models of iTILs and overall survival in 60 African American women with TNBC.

| Characteristic | Model 1 |  |  | Model 2* |  |  | Model 3* |  |  |
| --- | --- | --- | --- | --- | --- | --- | --- | --- | --- |
|  | HR | 95% CI | P-value | HR | 95% CI | P-value | HR | 95% CI | P-value |
| iTILs |  |  |  |  |  |  |  |  |  |
| 0% | 1.00 | --- |  | 1.00 | --- |  | 1.00 | --- |  |
| 1-10% | 0.32 | 0.12-0.90 | 0.031 | 0.42 | 0.11-1.64 | 0.21 | 0.34 | 0.06-1.89 | 0.22 |
| >10% | 0.18 | 0.05-0.67 | 9.9x10<sup>-3</sup> | 0.26 | 0.05-1.28 | 0.098 | 0.23 | 0.04-1.39 | 0.11 |
| Ordinal | 0.41 | 0.21-0.81 | 0.010 |  |  |  |  |  |  |
| Age (years) | 0.99 | 0.96-1.03 | 0.61 | 0.99 | 0.94-1.04 | 0.66 | 0.99 | 0.94-1.05 | 0.85 |
| Stage |  |  |  |  |  |  |  |  |  |
| Local | 1.00 | --- |  | 1.00 | --- |  | 1.00 | --- |  |
| Regional | 1.94 | 0.80-4.73 | 0.14 | 1.55 | 0.47-5.07 | 0.47 | 1.13 | 0.29-4.42 | 6.7x10<sup>-3</sup> |
| Distant | 4.60 | 1.08-19.6 | 0.039 | 24.5 | 1.97-304.3 | 0.013 | 39.6 | 2.78-565.5 | 0.86 |
| Chemotherapy sequence** |  |  |  |  |  |  |  |  |  |
| Adjuvant |  |  |  | 1.00 | --- |  | 1.00 | --- |  |
| Neoadjuvant |  |  |  | 3.31 | 0.85-13.0 | 0.086 | 3.29 | 0.74-14.6 | 0.12 |
| Surgery type |  |  |  |  |  |  |  |  |  |
| Breast conserving |  |  |  |  |  |  | 1.00 | --- |  |
| Mastectomy |  |  |  |  |  |  | 1.42 | 0.26-7.67 | 0.68 |
| Radiation |  |  |  |  |  |  |  |  |  |
| No |  |  |  |  |  |  | 1.00 | --- |  |
| Yes |  |  |  |  |  |  | 0.36 | 0.086-1.48 | 0.16 |

\* 21 observations deleted due to missing chemotherapy or type of surgery data.
\*\* Among patients with chemotherapy data.

### Gene expression patterns associated with TILs & survival

To better understand the molecular and cellular pathways—as represented by expression patterns—captured by histopathologically-defined iTILs in these TN tumors, we next interrogated ontologic pathway enrichment among transcripts associated with iTILs. At p<0.01, a total of 524 transcripts (326 coding, 198 non-coding) were associated with iTILs (**Table S2**). Among these, 224 transcripts were inversely associated with iTILs such that tumors with higher levels of iTILs had lower levels of expression of these transcripts. Of the 198 non-coding transcripts, 78 (36%) were long non-coding (lnc) RNAs, compared to only 26% of all noncoding transcripts on the array that correspond to lncRNAs. We evaluated functional pathway enrichment among the 326 significant coding transcripts using Ingenuity^®^ Pathway Analysis (IPA) software, finding that a wide range of immune cell types and pathways were represented, confirming that the significant transcripts from our analyses likely represent biologically relevant expression patterns. These fell largely into five general categories: (1) antigen presentation, (2) autoimmune signaling involving abnormal B cell recognition of self-antigens, (3) dendritic cell signaling, (4) B and T cell development and differentiation, and (5) T cell activation and signaling (**Table S3**).

While these 524 significant transcripts likely reflect the cell types encompassed by histopathologic TIL review, we sought to further refine this list to identify those also relevant to survival in TNBC. Thus, we next examined whether these 524 transcripts were associated with survival when accounting for iTIL levels, age and stage at diagnosis. A total of 34 of these transcripts were significantly associated with survival in this model (p<0.05) (Table 3), falling into two categories: (1) twelve transcripts that appeared to attenuate the effect of iTILs on survival and (2) twenty-two transcripts that appeared to enhance the effect of iTILs on survival.

**Table 3.**
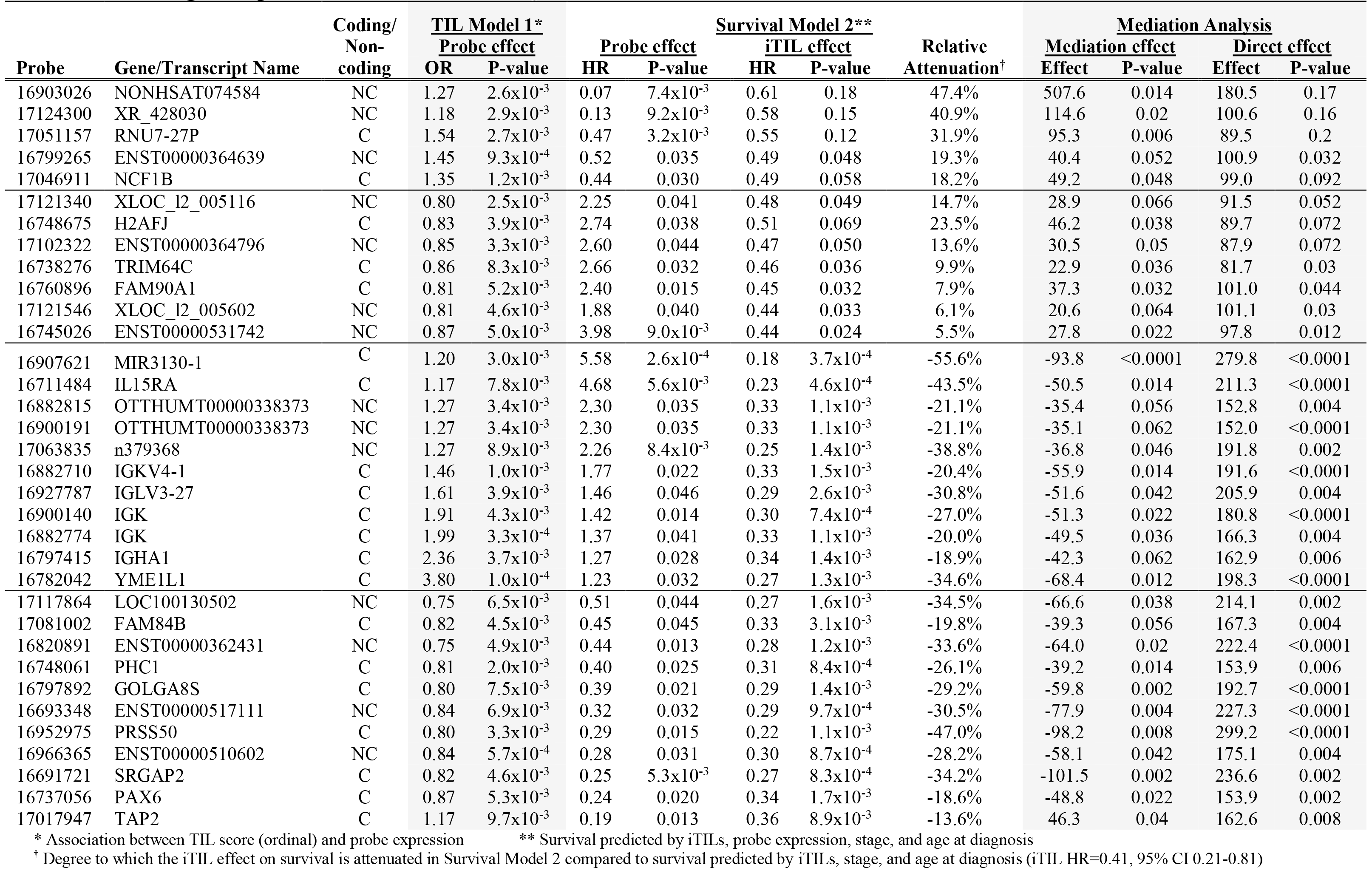
iTILs, gene expression and survival among 60 AA women with TNBC.

Within this first group of twelve transcripts, the magnitude of iTIL effect attenuation ranged from 5.5-47.4%, and formal mediation analysis indicated that all of these 12 transcripts may partially mediate the relationship between iTILs and survival with at least marginal statistical significance (Table 3). Interestingly, more than half of these transcripts were more highly expressed in tumors with lower levels of immune infiltrate, including *H2AFJ, TRIM64C*, and *FAM90A1*, suggesting that in a subset of TN tumors, iTILs may improve survival by suppressing the cells that highly express these transcripts. Within the second group of twenty-two transcripts, the effect of iTILs on survival strengthened by 13.6-55.6%. Half of these transcripts were more highly expressed in tumors with higher levels of immune infiltrate (e.g., *TAP2, PAX6, FAM84B, PHC1, GOLGA8S, SRGAP2, PRSS50*), but were themselves associated with worse survival, suggesting that the composition of iTILs in this subset of TN tumors may include a variety of immune cell types with contradictory effects on survival. The remaining half of these transcripts were more highly expressed in tumors with lower iTIL levels (e.g., *IGHA1, IGK, IGKV4-1, IGLV3-27, IGKV1OR2-1, YME1L1, IL15RA, MIR3130-1*), but were themselves associated with improved survival, suggesting that there may exist a subset of TN tumors where a compensatory mechanism is in place when immune response is limited. Overall, these patterns suggest that iTILs defined by histopathologic review of H&E slides capture multiple immune cell types with either competing or synergistic effects on survival.

### Immune cell subtypes

To tease apart the effects we observed in our agnostic gene expression analysis, we performed a cell type deconvolution analysis using the CIBERSORT program. There was a wide range in the distribution of cell types estimated to be present in these tumors using CIBERSORT. The most prevalent were naïve B cells (Mdn. 13.5%), CD8+ T cells (Mdn. 23.4%), naïve CD4+ T cells (Mdn. 8.9%), followed by resting NK cells (Mdn. 7.4%), monocytes (Mdn. 7.9%), and M2 macrophages (Mdn. 5.9%) (**Figure S3**). We required that at least 10% of tumors (n=6) had estimated proportions ≥1% for a given cell type to be included in subsequent analyses, resulting in exclusion of resting CD4+memory T cells, γδ-T cells, and neutrophils. We performed principal components analysis to evaluate the relationships between the remaining 19 immune cell types in these 60 TN tumors, finding that tumors clusters were inversely driven by the presence of CD8+ T cells and either naïve B cells or naïve CD4+ T cells. Further separation was influenced by the presence of either M1 or M2 macrophages (**Figure S4**). We evaluated the association between individual cell type proportions and survival adjusting for age, stage at diagnosis, and iTIL levels. Only naïve B cells were significantly associated with survival when considering immune cell types individually [Median: HR=2.43, 95% CI 1.07-5.55, p=0.035].

Due to the heterogeneity of these tumors with respect to patterns of immune infiltrate, evaluation of individual cell types might not capture underlying associations with survival. Thus, we next evaluated correlations between these immune cell subtypes and the 34 transcripts associated with iTILs and survival (Figure 2). Importantly, these 34 transcripts were not included among immune signature transcripts, indicating that expression of these 34 transcripts is independent of CIBERSORT cell proportion estimates. Associations between these 19 immune cell types and the 34 transcripts associated with iTILs and survival revealed striking patterns. First, transcripts that were expressed in the absence of iTILs and also associated with reduced survival were generally, though not exclusively, associated with decreased M1 macrophages, increased activated NK cells, and increased CD8+T cells. Second, transcripts that were expressed when iTILs were present but associated with reduced survival were generally associated with decreased monocytes, increased plasma cells, and increased naïve B cells. About half of these transcripts were also associated with reduced levels of resting mast cells, activated NK cells, and possibly CD8+ T cells. It is interesting to note that activated NK cells showed contradictory associations with transcripts associated with poor survival, dependent on whether those transcripts were also associated with the presence or absence of iTILs. Finally, transcripts associated with improved survival, regardless of the direction of their association with iTILs, had no clear patterns of association with immune cell types.

**Figure 2.**
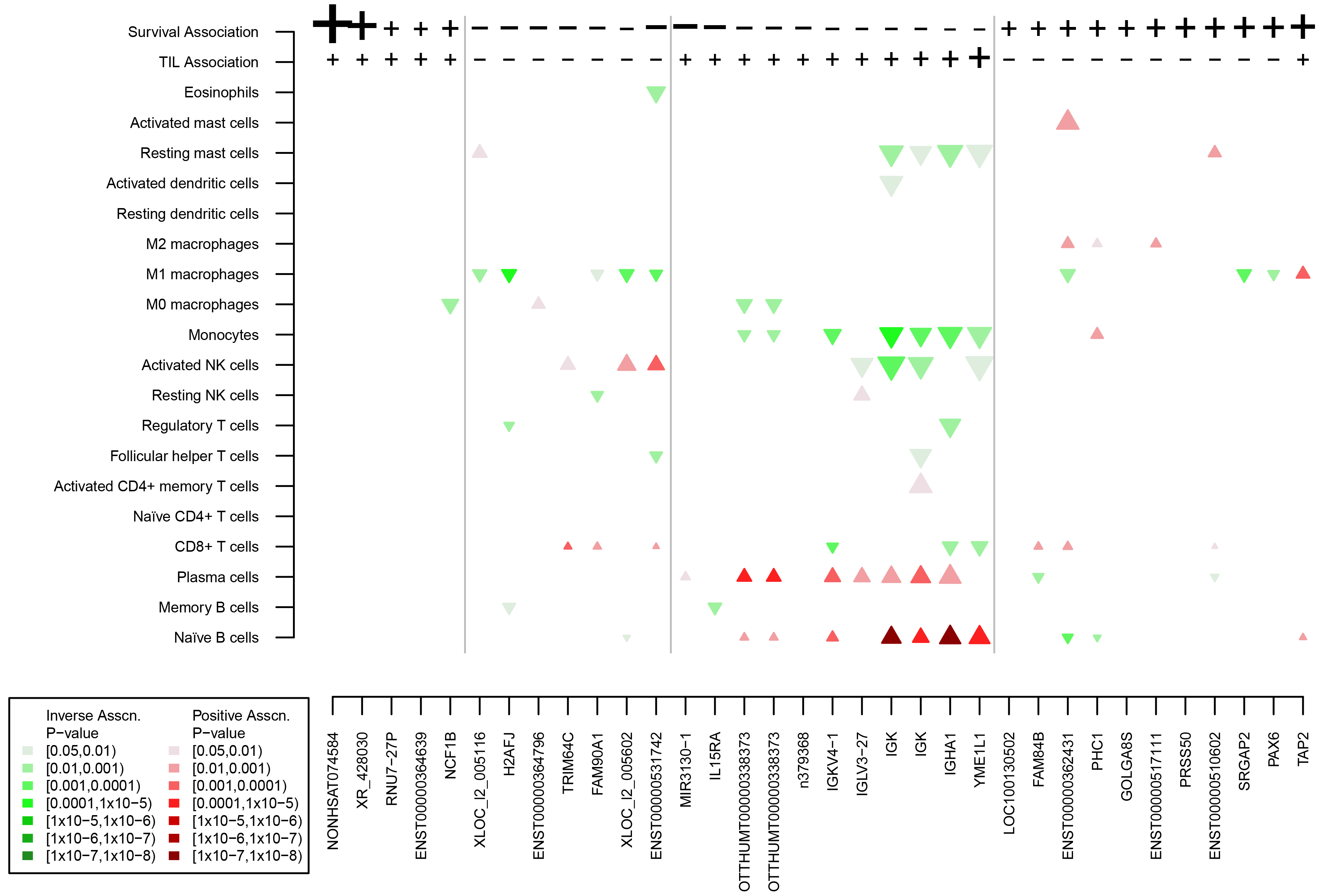
Immune cell type associations with 34 TIL– and survival-associated genes. Patterns of association between 19 immune cell subtypes (with at least 10% of tumors containing =1% cells) and the 34 genes found to be associated with both iTILs and survival are shown. Direction and strength of association between each gene and a) survival and b) iTILs are shown using “+” for positive associations or “−” for inverse associations, such that a “+” survival association indicates that increased expression of the gene is associated with better survival and a “+” iTIL association indicates that increased expression of the gene is associated with higher iTIL levels. Symbol size is proportional to the size of the effect estimate (see Table 3). Grey lines separate genes according to patterns of association with survival and iTILs. Triangles are shaded and oriented according to the direction of association between the cell type proportion and the gene of interest: downward blue=inverse association, upward red=positive association. Color intensity is proportional to statistical significance. Triangle size is proportional to log10 of the absolute value of the regression coefficient.

## DISCUSSION

Using the Metropolitan Detroit Cancer Surveillance System, our analysis of gene expression profiles from TN tumors among sixty African American women confirms that higher levels of immune infiltrate in tumors corresponds to improved survival. We found that transcripts significantly associated with both iTIL levels were implicated in pathways related antigen presentation, autoimmune signaling involving abnormal B cell recognition of self-antigens, dendritic cell signaling, B and T cell development and differentiation, and T cell activation and signaling, of which 34 were also associated with survival. These transcripts were associated with distinct immune cell subtype patterns involving M1 macrophages, activated NK cells, CD8+ T cells, naïve B cells, plasma cells, naïve B cells, and resting mast cells. These findings are novel in that our study is the first to focus on the association of immune cell levels and composition with respect to survival among African American women diagnosed with TNBC.

Two studies have been published investigating CIBERSORT estimates of 22 immune cell subtypes and clinical outcomes in breast cancer using publically available gene expression data (24,25). In the first, Ali et. al found that CD8+ T cells and activated memory T cells were associated with improved survival, while T regulatory cells, M0 macrophages, and M2 macrophages were associated with worse prognosis among ER-negative tumors (24). This study, however, was unable to distinguish between HER2-postive and HER2-negative tumors. Bense, et al. evaluated these cells type stratified by ER, PR, and HER2 status and found that higher fractions of neutrophils, resting mast cells, M2 macrophages, resting NK cells, and resting CD4+ memory T cells were associated with worse overall survival, and better overall survival was associated with higher fractions of plasma and naïve B cells among TNBC(25). Neither of these studies, however, considered race in their analyses. Our results provide evidence for conflicting roles for naïve B cells, plasma cells, resting mast cells, and novel roles for M1 macrophages, activated NK cells, monocytes in African American women with TNBC compared with these published studies.

Further, while we identified 16 transcripts associated with improved survival and iTILs, we did not identify any clear immune cell subtype patterns associated with improved survival. Several of these 16 transcripts (*FAM84B, PAX6, PHC1, SRGAP2*) have evidence for a role in breast cancer clinical outcomes, both *in vitro* and *in vivo*, including TNBC. *FAM84B*, also known as Breast cancer-membrane associated protein 101, is located in the chromosome 8q24 region associated with susceptibility to breast, prostate, colon and other cancers (35,36). FAM84B was identified as uniquely expressed in breast cancer cell membranes compared to normal breast epithelial cells (37), and upregulation was also associated with progression and pathological response to neoadjuvant chemoradiation in esophageal squamous cell carcinoma (38, 39). *PAX6* (Paired box 6) encodes a homeobox and paired domain-containing DNA-binding protein and transcription factor. Hypermethylation of *PAX6* was associated with hormone receptor positive compared to hormone receptor negative breast cancer, and *PAX6* overexpression was found in 43.9% of ER-negative breast cancers and was associated with worse survival independent of subtype (40–42). *PHC1* (Polyhomeotic Homolog 1) encodes a component of a protein complex required to maintain the transcriptionally repressive state of many developmental transcripts, and was utlilized in an MCM2-targeted treatment strategy that successfully induced DNA damage and subsequent apoptosis in murine invasive breast cancer cells (43). Finally, *SRGAP2* (SLIT-ROBO Rho GTPase Activating Protein 2) encodes a protein that regulates actin dynamics to regulate cell migration and differentiation and plays a role in cortical neuron development, and was found to be upregulated in early stage recurrent TNBC (44).

Compared to other breast cancer subtypes, patients diagnosed with TNBC have higher risk of recurrence and distant metastases (33,45–47). Given the lack of expression of hormone receptors and HER2 amplification, primary treatment options for TNBC are limited to surgery and systemic therapies (48). Indeed, in our cohort of women who received surgical treatment for breast cancer, we noted that the majority underwent a mastectomy and received chemotherapy. While previous reports have suggested that there are no significant differences in local control or survival between type of operative therapy (breast-conserving surgery versus mastectomy) for patients (49), TILs were established as predictive markers of patient response to neoadjuvant chemotherapy in clinical trials (28,30,31).

Pathological assessment of TILs and tumors and use of the Metropolitan Detroit Cancer Surveillance System to identify cases of TNBC among African American women allowed for detailed patient follow-up. While TILs have been shown to be predictive of treatment response in breast cancer (30, 31), we were unable to collect detailed information on chemotherapy regimens, which would have provided an assessment on the predictive value of TILs on treatment response in the African American population. Use of the Affymetrix platform for gene expression profiling of FFPE samples and data processing allowed for accurate interpretation of our findings (50). We acknowledge that a larger sample size for expression profiling of TNBC tumors from African American women is needed to further examine the association as well as the prognostic value of novel genetic mediators identified in this study associated with TILs and survival.

Our findings in this cohort of African American women with TNCB establish novel roles for several immune cell types in AAW with TNBC compared to published data from largely European white populations. We also identified sets of transcripts associated withimproved survival that had no clear pattern of association with immune cell types, indicating that these transcripts may be biomarkers for tumor cell response to immune infiltration. Future studies must be conducted to validate these findings and identify determinants of these patterns of immune response in TNBC, including genetic, hormonal, and health-related factors.

